# Explainable machine learning identifies candidate shared neuroanatomical features in Alzheimer’s and Parkinson’s via importance inversion transfer

**DOI:** 10.64898/2026.04.10.717735

**Authors:** Daniele Caligiore, Simone Torsello, the Alzheimer’s Disease Neuroimaging Initiative

**Author notes:** A complete listing of ADNI investigators can be found at: http://adni.loni.usc.edu/wp-content/uploads/how_to_apply/ADNI_Acknowledgement_List.pdf.

## Abstract

Despite significant neurobiological and pathological overlaps, Alzheimer’s and Parkinson’s diseases—the primary threats to healthy aging—are still managed as distinct clinical entities. Standard machine learning exacerbates this diagnostic fragmentation by prioritizing divergent markers over shared traits, thereby obscuring the invariant foundations of neurodegeneration. This study introduces Importance Inversion Transfer, an explainable machine learning framework designed to identify neuroanatomical invariants across the neurodegenerative spectrum. Prioritizing structural stability over discriminative utility isolates a shared pathological core consisting of ten regional volumetric anchors, validated through an inductive protocol with high diagnostic fidelity (AUC = 0.894). The identified morphological continuum between healthy aging and neurodegeneration delineates shared structural substrates consistent with—though not demonstrative of—a potential common early-phase vulnerability. Aligned with the Neurodegenerative Elderly Syndrome hypothesis, this evidence establishes a possible paradigm for early, system-level diagnosis.

## 1 Introduction

Alzheimer’s disease (AD) and Parkinson’s disease (PD) constitute the foremost causes of neurodegeneration in geriatric populations, imposing a mounting strain on global healthcare infrastructure [1, 2]. Conventional clinical paradigms categorize these disorders as separate entities defined by symptomatic silos: while AD manifests primarily through progressive cognitive erosion and memory loss, PD is clinically distinguished by hallmark motor impairments, including bradykinesia, tremor, and rigidity [3–5]. However, emerging evidence highlights substantial commonalities at the molecular and cellular levels which frequently manifest decades prior to clinical onset [6–11].

These convergences have led to the hypothesis that AD and PD may share partially overlapping disease mechanisms, potentially reflecting a common early vulnerability phase. This perspective has been formalized as the Neurodegenerative Elderly Syndrome (NES) hypothesis [8, 12]. The NES framework delineates a three-phase progression: an initial *seeding stage* defined by subclinical *α*Syn and monoaminergic failure; a *compensatory stage* wherein homeostatic regulation preserves functional integrity despite incipient decay; and a terminal *bifurcation stage* in which damage to specific neural circuits yields phenotype-specific manifestations. Despite this theoretical foundation, mapping the shared neuroanatomical backbone that sustains the syndrome prior to clinical divergence remains a critical challenge in geriatric neurology.

Exploring the plausibility of the NES hypothesis requires analytical methods capable of *capturing both shared and divergent features* within complex, multimodal datasets. While machine learning (ML) offers powerful pattern detection capabilities, clinical application demands transparency to ensure safety and reliability. *Explainable machine learning (XML)* addresses this requirement by identifying the specific features driving model predictions, thereby enabling a mechanistic interpretation of neurodegenerative trajectories. Recent literature underscores the versatility of XML in this domain, spanning from broad applications in diagnosis and treatment monitoring [13] to the identification of manifestations through digital phenotyping [10] and the optimization of multi-target drug combinations [14]. Specifically, XML has proved pivotal in unraveling sex-specific pathological signatures in both PD and AD, facilitating the development of clinician-tailored diagnostic interfaces [15, 16]. However, despite these advancements, current diagnostic models and XML protocols primarily emphasize feature selection based on maximum divergence. This heavy reliance on discriminative utility inherently obscures the invariant structural anchors that constitute the shared pathological core of neurodegeneration.

To address this limitation, the present study employs the Importance Inversion Transfer (IIT) mechanism [17] to map the set of candidate invariant neuroanatomical features of AD and PD. Drawing on the identification of general invariant principles within complex multiscale systems [18], the IIT logic inverts standard selection protocols by prioritizing neuroanatomical descriptors that exhibit high structural invariance and statistical stability across disease states. This methodological shift prioritizes features with low discriminative power, enabling the identification of candidate regions that may reflect shared structural properties across disease states.

Analysis of multi-regional brain volumes across Healthy Controls (CN), PD, and AD cohorts suggests the potential existence of a unified neuroanatomical signature. The identification of stable structural anchors points toward a common pathological trunk, encouraging a transition from phenotype-specific classification toward a broader system-level view of brain aging [19, 20]. Such findings offer a preliminary framework for intervention strategies aimed at shared neurodegenerative mechanisms during stages preceding definitive clinical bifurcation.

## 2 Results

### 2.1 Volumetric Signatures and Phenotypic Bifurcation

Morphometric profiling across 84 brain regions delineates distinct structural signatures for AD and PD relative to healthy aging (Table 5). Analysis reveals patterns consistent with an antagonistic relationship: AD manifests as a systemic structural collapse of limbic and associative networks, whereas PD exhibits localized shifts and selective volumetric expansion.

AD pathology shows aggressive progression centered on the medial temporal lobe. Entorhinal (LH: 34.11%) and amygdala (LH: 32.21%) atrophy identify the primary sites of tissue loss (*p <* 0.001), followed by significant hippocampal contraction (LH: 28.36%, *p <* 0.001). This neuroanatomical degradation extends symmetrically to the inferior temporal gyrus and the precuneus. Widespread tissue depletion coincides with bilateral expansion of the lateral ventricles (*>* 85%) and total CSF volume (34.07%), marking the macroscopic stage of advanced neurodegeneration.

The PD cohort displays a divergent morphological trajectory, frequently maintaining statistical contiguity with the healthy baseline. Significant volumetric expansion occurs in the brain stem (5.25% increase, *p* = 0.030) and the hemispheric cerebral white matter (LH: 4.92%; RH: 5.07%, *p <* 0.05). A similar hypertrophic trend appears in the left choroid plexus, where expansion reaches 10.40% without meeting the conventional significance threshold (*p* = 0.060).

Volumetric distributions identify a bifurcation in neurodegenerative paths between AD and PD. While AD manifests through moderate white matter contraction (9– 10%), PD sustains or increases white matter volume. Similarly, subcortical nuclei, including the thalamus and putamen, show significant volumetric separation from the healthy baseline in AD, yet remain statistically indistinguishable from controls in the PD cohort.

### 2.2 Classification Performance and Diagnostic Benchmarking

Performance benchmarking of three ML architectures, *Random Forest* (RF), *Gradient Boosting* (GB), and *Logistic Regression* (LR), through a multi-seed inductive validation protocol identifies the diagnostic potential of regional brain volumetric signatures. As summarized in Table 1, the RF classifier demonstrates superior diagnostic power for multi-cohort separation, yielding a mean accuracy of 73.2% and a robust Area Under the Curve (AUC) of 0.895.

**Table 1.** Global 3-way classification performance (Healthy vs. PD vs. AD). Metrics represent ensemble averages marginalized over 10 independent random seeds. Accuracy, F1-macro, and ROC-AUC scores confirm the robustness of volumetric features for primary cohort identification.

| Model | Accuracy | F1-macro | ROC-AUC |
| --- | --- | --- | --- |
| **RF** | 0.732 | 0.730 | 0.895 |
| GB | 0.709 | 0.707 | 0.878 |
| LR | 0.685 | 0.686 | 0.849 |

To isolate the set of candidate invariant neuroanatomical features of AD and PD, the importance inversion logic was applied to a binary disease-only classification manifold. By removing CN from the discovery phase, the models were forced to prioritize structural descriptors with the lowest discriminative utility specifically between the two pathologies. This strategy ensures that the identified structural anchors represent the purest invariants of the neurodegenerative seeding phase, effectively filtering out any signal related to the general transition from healthy aging to disease. The binary diagnostic challenge specifically targeting the differentiation between PD and AD reveals an even higher degree of separability (Table 2). In this task, GB achieves a peak accuracy of 93.8% and an AUC of 0.981. This performance jump indicates that while the global transition from healthy aging is complex, the phenotype-specific signatures that distinguish PD from AD are highly unambiguous once the healthy baseline variance is removed.

**Table 2.**
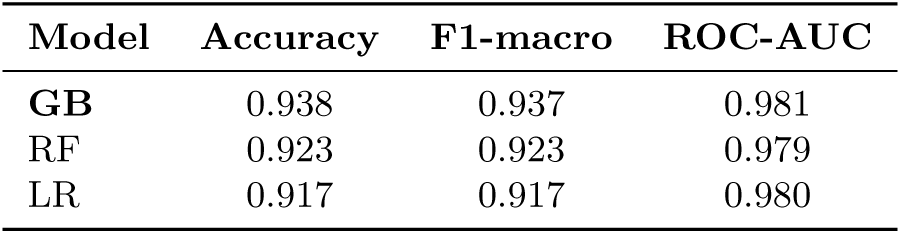
Binary classification performance (PD vs. AD bifurcation). Summary of performance metrics for the direct comparison between disease phenotypes. High accuracy and AUC values across all models indicate nearly absolute structural differentiation between PD and AD cohorts.

More in details, Figure 1 reveals distinct structural regimes. In the global 3-way task, the AD cohort reaches a classification accuracy of 88.6%, suggesting a highly specific structural fingerprint. Conversely, the significant overlap between CN and PD—where 33.3% of PD subjects are misclassified as CN—highlights a morphological continuum characteristic of the early neurodegenerative seeding phase.

**Fig. 1.**
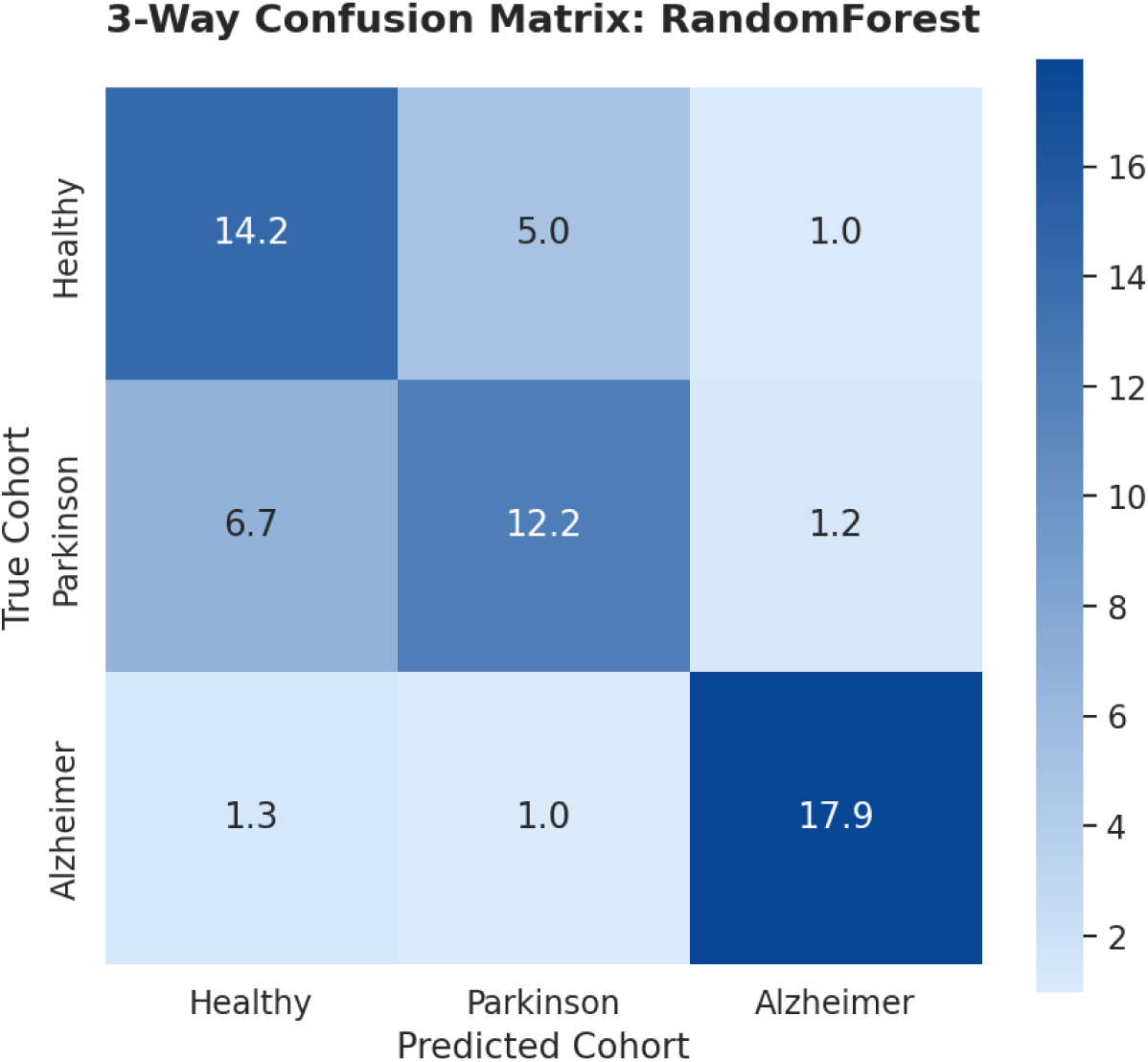
Global cohort classification. Average confusion matrix marginalized over 10 independent random seeds for the 3-way RF classifier. Values represent the mean number of subjects per test fold (*n_total_* = 60.5 per fold). The high accuracy for AD (88.6%) contrasts with the PD/Healthy overlap (33.3%), highlighting the early-stage morphological continuum.

This statistical proximity between PD and healthy aging indicates that volumetric shifts in early PD often reside within the variance of normal physiological aging. This phenomenon prevents total separation of the manifolds until the bifurcation stage is reached, further justifying the implementation of the IIT logic (see next section). By bypassing the classificatory noise inherent in these overlapping distributions, the IIT mechanism successfully isolates the stable structural anchors that remain invariant across the neurodegenerative spectrum.

The diagnostic efficacy of the binary task is further elucidated by the average confusion matrix for the GB classifier (Figure 2). The results suggest a strong separation between the two pathological states, with a mean of 18.8 PD subjects (93.1%) and 19.1 AD subjects (94.6%) correctly identified per test fold. The marginal misclassification rate—averaging only 1.4 PD subjects identified as AD and 1.1 AD subjects identified as PD—suggests that, once the healthy baseline variance is removed, the neuroanatomical footprints of AD and PD occupy mutually exclusive morphometric manifolds. This high categorical distinctness provides quantitative evidence for the phenotypic bifurcation stage of the NES. While the seeding phase is characterized by structural contiguity, the bifurcation stage is defined by the emergence of non-overlapping volumetric signatures.

**Fig. 2.**
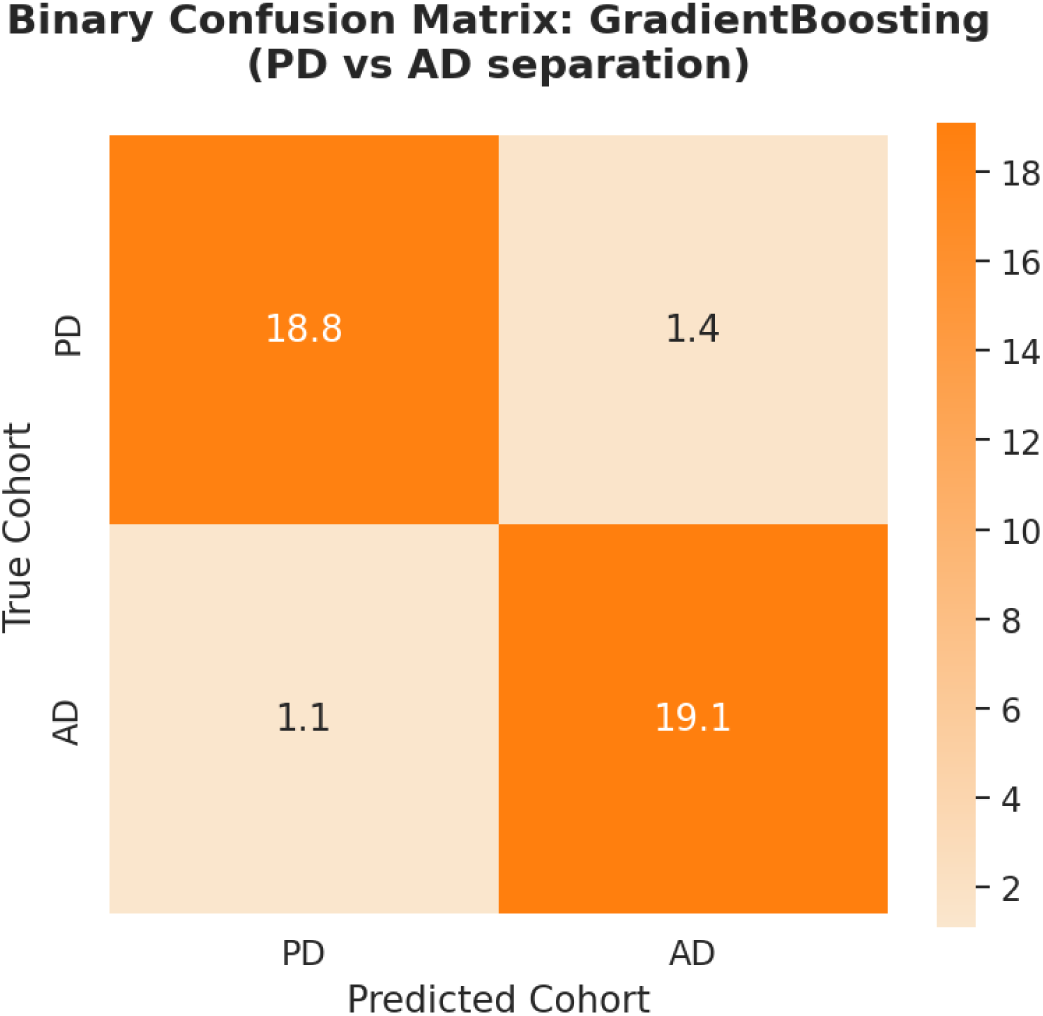
Binary AD vs. PD separation. Average confusion matrix for the GB classifier in the binary task. The diagonal dominance (93.8% overall accuracy) indicates that AD and PD occupy distinct neuroanatomical manifolds when compared directly, facilitating precise phenotypic bifurcation.

### 2.3 Diagnostic Separators and Structural Anchors

Borda consensus ranking in the binary classifiers highlights the brain regions that most strongly drive phenotypic separation (Fig. 3). Structural profiling reveals that the *LH Hippocampus*, *RH Amygdala*, *LH Entorhinal cortex*, and *RH Hippocampus* exhibit the highest discriminative utility. These areas, characterized by extreme statistical separation (*p <* 0.001), define the bifurcation stage of the neurodegenerative process.

**Fig. 3.**
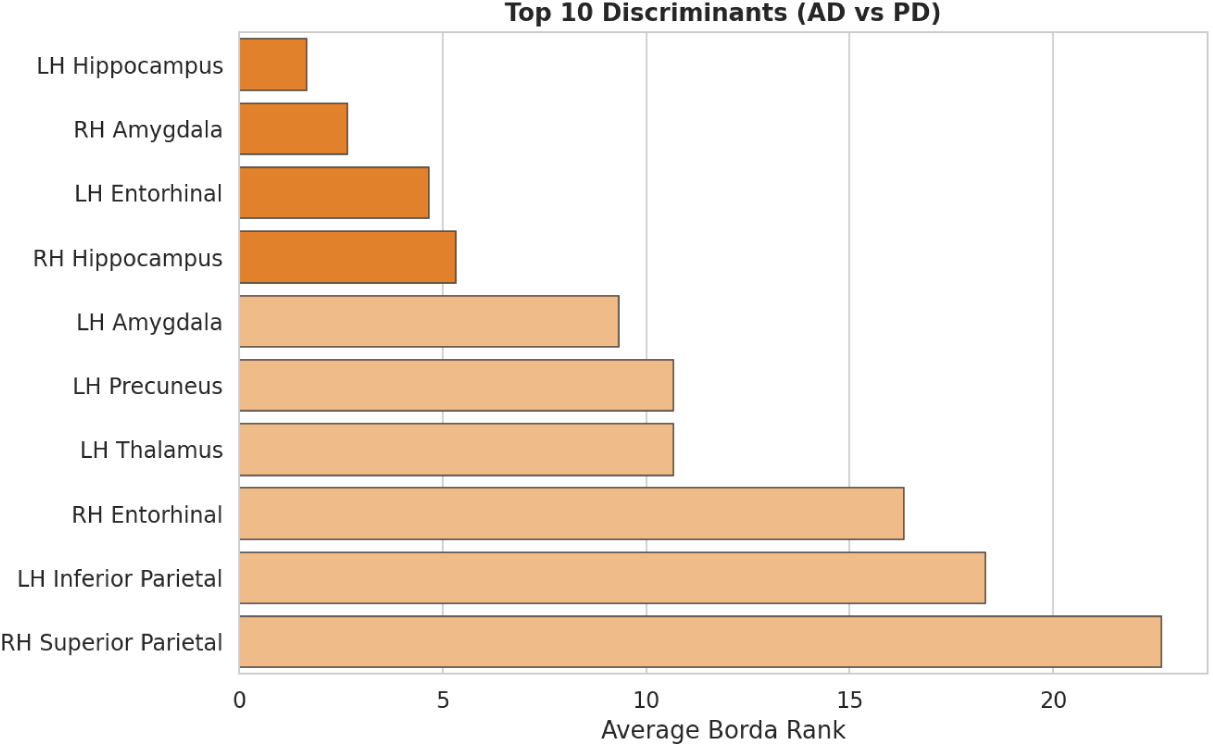
Top Classificatory Discriminants. Borda consensus ranking identifying the brain regions with maximum discriminative utility between AD and PD (Bifurcation markers).

Conversely, the IIT score isolates the structural anchors that constitute the shared pathological core (Fig. 4). The *LH Transverse Temporal* cortex and the *RH Choroid Plexus* emerge as the top-ranked features, identifying a parsimonious neuroanatomical backbone that remains synchronized across pathologies. Notably, the *RH Amygdala* represents the sole descriptor present in both top-tier rankings. Although it serves as a primary driver for diagnostic separation (ranking 2nd in Fig. 3), it simultaneously functions as a structural anchor (Fig. 4). The Discussion section further examines the pathophysiological basis of this dual positioning, identifying the region as a critical hybrid node within the neurodegenerative manifold.

**Fig. 4.**
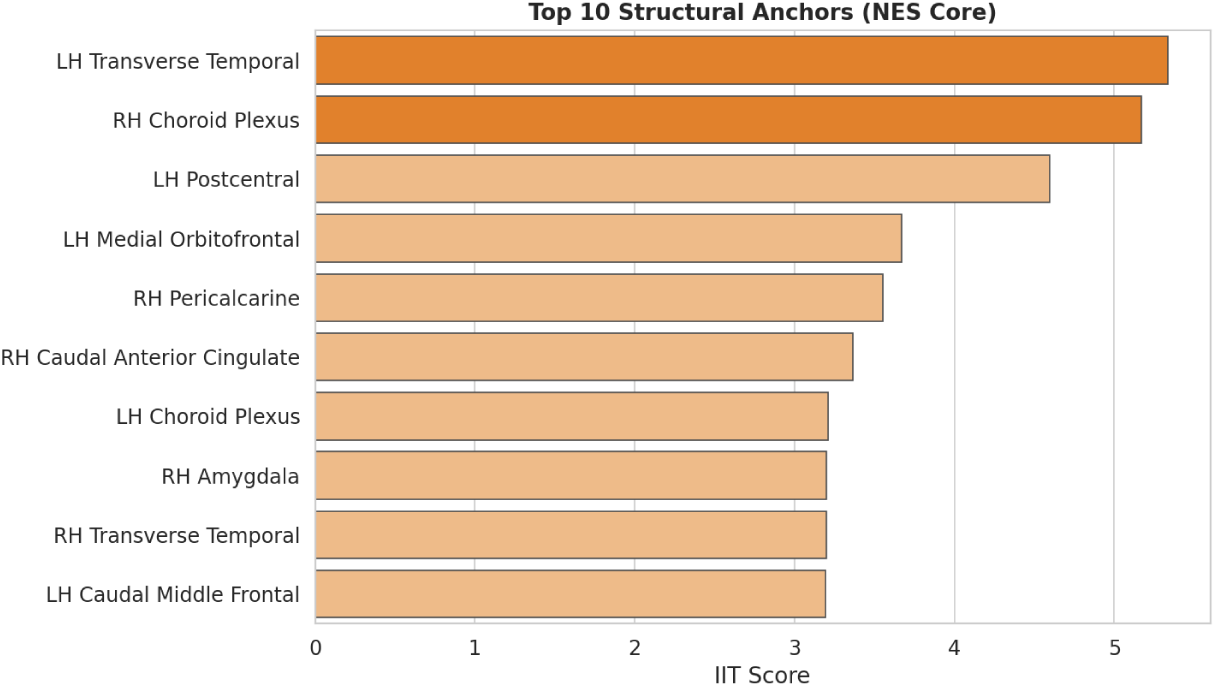
Structural Anchors and the NES Core. IIT score ranking identifying the regions with maximum pathological invariance across the neurodegenerative spectrum.

### 2.4 Hierarchical Audit of the Neuroanatomical Backbone

Statistical validation of top-ranked regions highlights a distinct hierarchy within the shared neuroanatomical core (Table 3). The *LH Transverse Temporal* cortex (*p* = 0.1063) and *RH Choroid Plexus* (*p* = 0.0510) function as *Master Anchors*. These regions show reduced statistical separation between AD and PD, indicating limited discriminatory power in this dataset. Minimal effect sizes (*<* 0.34) further establish these nodes as the invariant neuroanatomical locus of the NES seeding stage.

**Table 3.** Independent Statistical Validation of the NES Core Hierarchy. Regions are ranked by their Binary IIT Score. Master anchors (in bold) exhibit phenotypic invariance between AD and PD (*p_PD/AD_* ≥ 0.05). Transition pillars represent regions with high structural centrality where pathological trajectories begin to diverge significantly (*p_PD/AD_ <* 0.05).

| Brain Region | IIT Score | $p_{PD/AD}$ | Effect Size |
| --- | --- | --- | --- |
| **LH Transverse Temporal** | 5.4650 | 0.1063 | 0.2860 |
| **RH Choroid Plexus** | 5.4435 | 0.0510 | 0.3358 |
| LH Postcentral | 4.6012 | 0.0000 | 0.8847 |
| LH Medial Orbitofrontal | 3.7156 | 0.0000 | 1.1178 |
| RH Pericalcarine | 3.5960 | 0.0001 | 0.6292 |
| RH Caudal Ant. Cingulate | 3.4241 | 0.0047 | 0.5306 |
| LH Choroid Plexus | 3.2874 | 0.0037 | 0.4120 |
| RH Amygdala | 3.2249 | 0.0000 | 1.6916 |
| RH Transverse Temporal | 3.2115 | 0.0177 | 0.3540 |
| LH Caudal Mid. Frontal | 3.2084 | 0.0000 | 0.8642 |

The remaining structures—including the *LH Postcentral* gyrus, *LH Medial Orbitofrontal* cortex, and *RH Amygdala*—constitute *Transition Pillars*. Despite significant volumetric divergence in magnitude (*p <* 0.01; Effect Size *>* 0.53), these regions maintain absolute rank-order consistency (*ρ* = 1.0). This dissociation between damage magnitude (Effect Size) and distributional pattern (Spearman’s *ρ*) suggests a synchronized pathological program across the structural backbone. Specifically, the high effect sizes in the *RH Amygdala* (1.6916) and *LH Medial Orbitofrontal* cortex (1.1178) mark the onset of a bifurcation stage where shared seeding yields to phenotype-specific atrophy trajectories. Uniform Spearman correlations (*ρ* = 1.0) across all investigated nodes confirm a perfectly preserved structural hierarchy. While atrophy magnitude and phenotypic divergence (*p_P D/AD_*) vary, the invariant rank-order of these anchors suggests them as stable topographical landmarks within the neurodegenerative landscape.

### 2.5 Bifurcation and Invariance in Neuroanatomical Manifolds

Statistical profiling of regional brain volumes delineates two distinct morphometric regimes: phenotypic bifurcation (Fig. 5) and pathological invariance (Fig. 6).

**Fig. 5.**
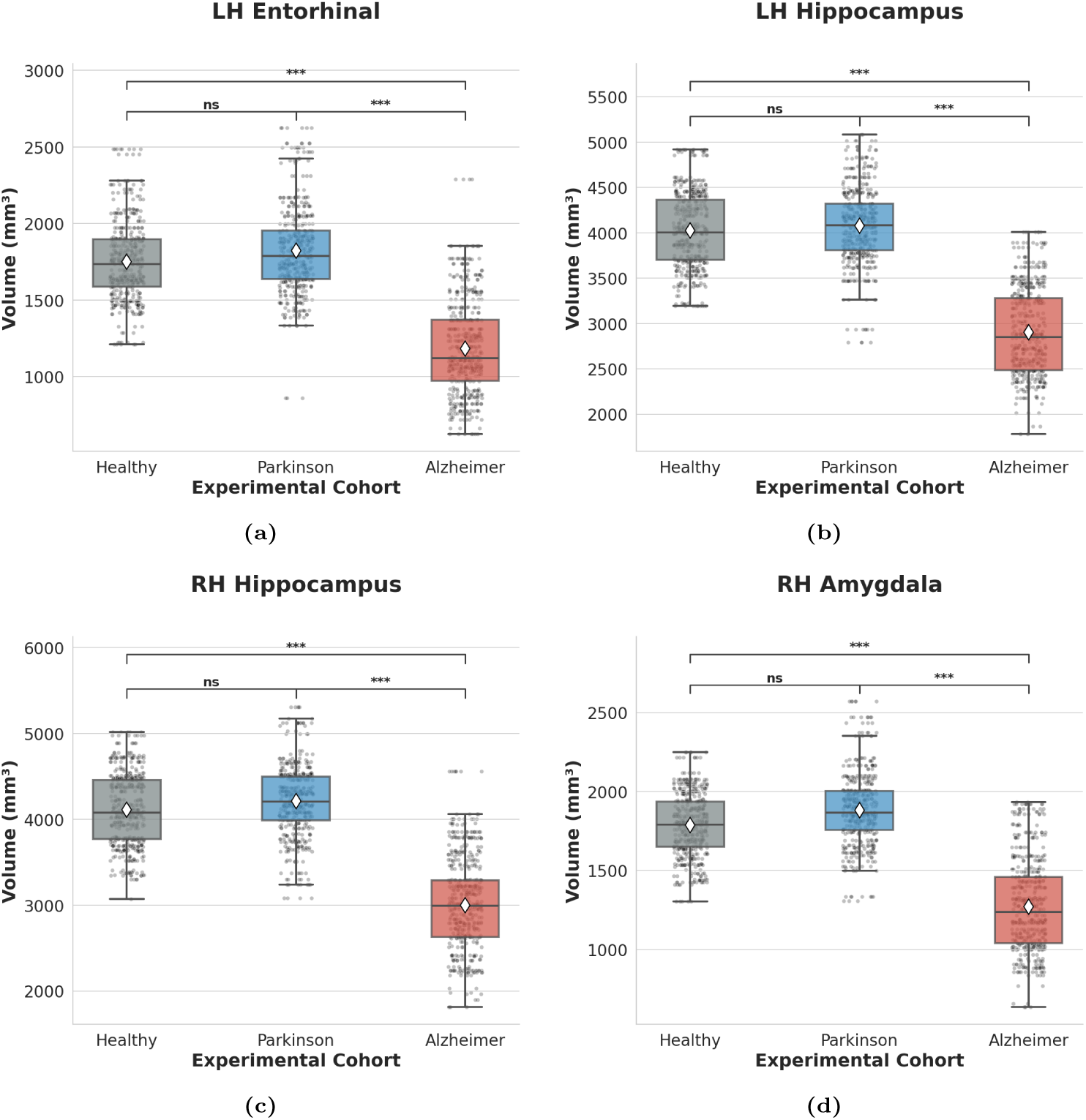
Morphometric markers of phenotypic bifurcation. (**a–d**) *LH Entorhinal* cortex, bilateral *Hippocampus*, and *RH Amygdala* showing extreme volumetric separation (*p <* 0.001) between AD and PD cohorts. These regions define the stage where phenotype-specific trajectories diverge from the common backbone. White diamonds indicate ensemble means; individual data points illustrate bootstrapped variance. Significance brackets denote post-hoc Dunn’s test results (∗ ∗ ∗*p <* 0.001, ns: not significant).

**Fig. 6.**
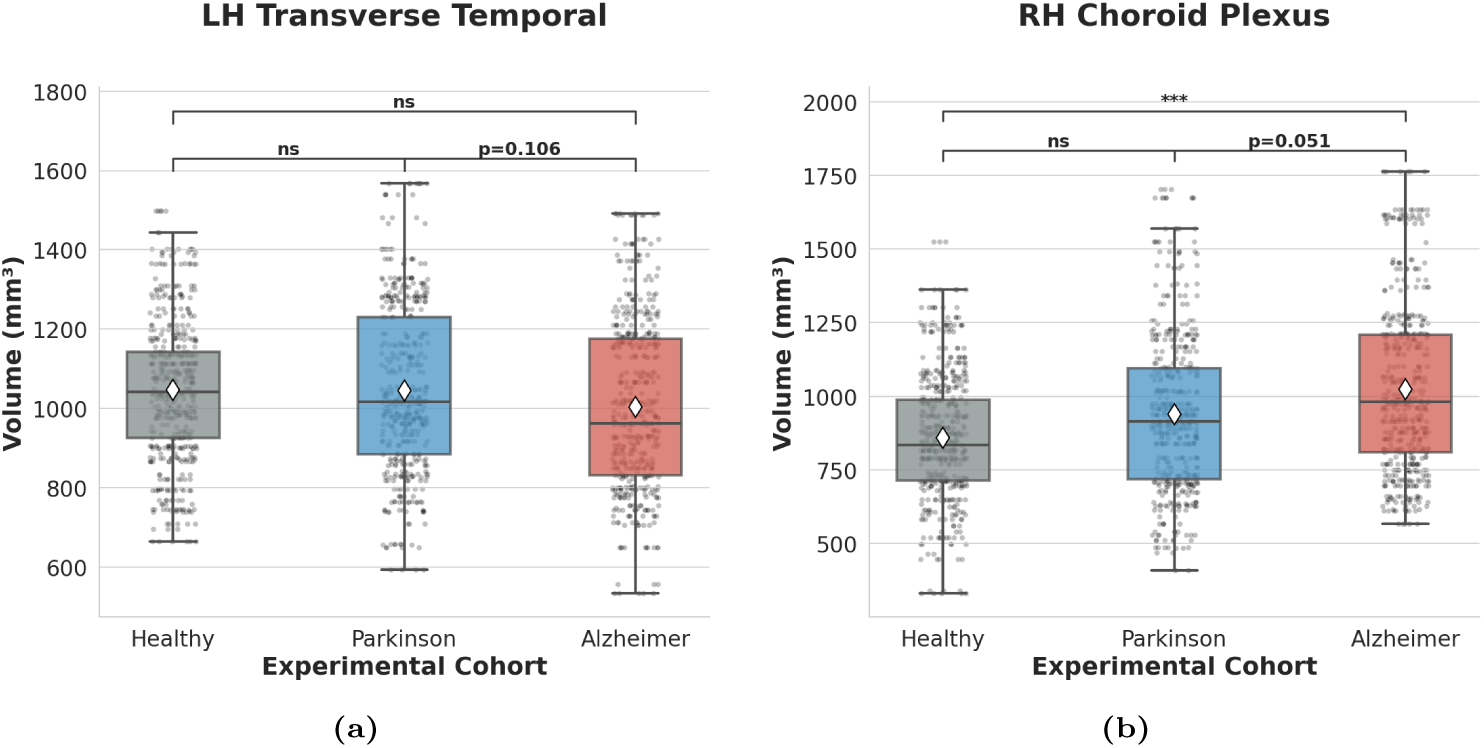
Structural Master Anchors of the neurodegenerative backbone. (**a,b**) *LH Transverse Temporal* cortex and *RH Choroid Plexus* exhibiting statistical indistinguishability (*p* ≥ 0.05) between AD and PD cohorts. These nodes identify the invariant seeding phase of the syndrome. White diamonds indicate ensemble means; individual data points illustrate bootstrapped variance. Significance brackets denote post-hoc Dunn’s test results (ns: not significant).

Ensemble distributions derived from 10 independent stochastic realizations visualize the transition from invariant seeding to symptomatic differentiation across six representative regions. Primary discriminants—specifically the *LH Entorhinal* cortex, the bilateral *Hippocampus*, and the *RH Amygdala*—delineate a clear structural divergence between the two neurodegenerative trajectories (Fig. 5a–d). In these hubs, the AD cohort exhibits profound volumetric collapse relative to both CN and the PD group (*p <* 0.001). Conversely, the PD cohort maintains statistical contiguity with the healthy baseline (*ns*), establishing these regions as the definitive markers of the bifurcation stage.

In contrast, structural anchors identify a shared neuroanatomical core where AD and PD trajectories remain synchronized. The *LH Transverse Temporal* cortex (Fig. 6a) demonstrates global stability across cohorts, with no significant difference between the pathological groups (*p* = 0.106).

The *RH Choroid Plexus* (Fig. 6b) functions as a critical transition node within this shared backbone. While AD subjects exhibit significant expansion relative to controls (*p <* 0.001), the direct comparison between AD and PD yields statistical indistinguishability (*p* = 0.051). This hierarchical significance suggests a shared hypertrophic signature that manifests fully in the AD phenotype while remaining in a latent, compensatory stage in PD (*ns* vs. Healthy). Such statistical proximity, despite significant deviation from the healthy baseline in the AD group, indicates the existence of a shared neuroanatomical backbone that characterizes the seeding phase of the neurodegenerative spectrum.

## 3 Discussion

### 3.1 Divergent Morphological Signatures and Pathological Trajectories

Regional volumetric profiling across 84 brain regions identifies distinct structural manifolds for healthy aging, PD, and AD (Table 5). Aggressive atrophy in the AD cohort perfectly reproduces established signatures of medial temporal and temporo-parietal decay [21]. The entorhinal cortex and hippocampus exhibit profound volumetric contraction, reflecting the initial stages of tau-related neurodegeneration within the medial temporal memory system [22–24]. This coordinated decay extends through limbic circuits to the amygdala and systemic hubs of the default mode network, such as the precuneus and temporal association areas [25–28]. Systemic tissue depletion culminate in a dramatic bilateral expansion of the lateral ventricles (*>* 80%), serving as a macroscopic surrogate for global volume loss and cognitive deterioration [24, 29].

In contrast, the PD cohort exhibits relative structural preservation and prominent instances of “negative atrophy.” Expansion in the brain stem and cerebral white matter (+4.92%) challenges traditional models of Parkinsonian progression, suggesting that regional volumetric increases represent a hallmark of the early-to-mid stage disease manifold [30–33]. Biologically, this phenomenon likely reflects the impact of *neuroinflammation* and reactive gliosis. The accumulation of *α*-synuclein triggers microglial activation and astrogliosis, resulting in a detectable increase in regional volumes that effectively masks concurrent neuronal depletion [34–38]. Statistical contiguity with healthy baselines persists in PD even alongside profound functional connectivity deficits, identifying a state of structural resilience or latency absent in the AD phenotype [39].

These divergent signatures suggest a fundamental pathophysiological distinction. While AD manifests as a systemic structural collapse, PD follows a topographically localized trajectory driven by a system-level failure in brain homeostasis [6, 40, 41]. Bilateral hypertrophy of the Choroid Plexus provides key evidence for this global disruption. As the primary site for cerebrospinal fluid production and waste clearance, expansion of the Choroid Plexus highlights a compensatory attempt to manage brainwide proteinopathic overload. This failure of the brain “drainage system” suggest that systemic neuro-inflammatory responses and impaired glymphatic clearance may effectively mask neuronal loss during early stages [42, 43].

Furthermore, brain stem and subcortical expansion suggests compensatory self-reorganization. Mirroring the neural remodeling observed in the thalamus and middle temporal gyrus of PD patients, these volumetric shifts likely reflect enhanced sensorimotor interactions or adaptive neural remodeling in response to progressing pathology [44]. Such dynamic subcortical morphology often correlates with clinical status and cognitive resilience [45]. Antagonistic white matter trajectories—expansion in PD (+4.92%) versus collapse in AD (-9.78%)—mark a definitive bifurcation in neurodegenerative paths. PD-specific neuro-inflammatory remodeling contrasts with the irreversible axonal loss characterizing AD [6, 46, 47].

The antagonistic trajectories identified between AD and PD provide significant diagnostic utility, establishing that neuroanatomical divergence precedes the point where phenotypic expressions reach diagnostic maturity. The relative stability of subcortical nuclei in PD, contrasted with their systemic decay in AD, suggests robust structural signatures for differential screening. Crucially, the implementation of rigorous normalization protocols ensures that these findings—specifically the observed volumetric resilience and “negative atrophy” in the PD cohort—reflect genuine biological phenomena rather than methodological scaling artifacts. By accounting for individual differences in total intracranial volume and demographic variance, the analysis captures a critical transition zone of biological resilience unique to the PD phenotype. Consequently, these results are compatible with the hypothesis that AD and PD may share early structural features before diverging into disease-specific trajectories. Early systemic failure in AD contrasts with a prolonged phase of homeostatic struggle in PD, supporting the hierarchical logic of the NES backbone.

### 3.2 Compensatory Latency and Phenotypic Bifurcation

High-accuracy binary classification (*AUC* = 0.981, Table 2) suggests that AD and PD occupy mutually exclusive morphometric manifolds during advanced disease stages. Yet, the global 3-way task reveals a critical asymmetry: while the AD signature remains highly distinct (88.6% accuracy), the significant misclassification of PD subjects as Healthy (33.3%) indicates the morphological continuum underlying early neurodegeneration (Fig. 1). This statistical proximity is consistent with the proposed compensatory stage within the NES framework, wherein homeostatic mechanisms effectively mask structural decay within the variance inherent in physiological aging.

As discussed in the context of antagonistic trajectories (cf., Section 3.1), the observed “negative atrophy” in the PD brain stem and cerebral white matter highlights the macrostructural footprint of reactive plasticity or neuro-inflammatory upregulation. This hypertrophic signature suggests a systemic effort to maintain network stability, potentially driven by the upregulation of noradrenergic and serotonergic systems in response to early dopaminergic depletion [8].

The pronounced contrast between this structural resilience in PD and the aggressive limbic collapse characterizing AD necessitates the application of IIT logic. By isolating invariant structural anchors that persist across the AD–PD transition, the framework captures latent pathological signatures well before clinical phenotypes achieve diagnostic maturity. This non-linear progression, where compensatory remodeling delays the emergence of hallmark symptoms, supports homeostatic markers as primary diagnostic indicators capable of bypassing the classificatory noise inherent in the bifurcation stage.

### 3.3 Anatomical Divergence and the NES Core Hierarchy

The quantitative divergence between Borda-driven separators and IIT-driven anchors delineates the structural transition from shared pathological seeding to phenotypic bifurcation (Figures 3, 4). Unlike traditional diagnostic paradigms prioritizing the *Hippocampus* for classification accuracy, IIT isolates the *RH Choroid Plexus* and the *LH Transverse Temporal* cortex as invariant master anchors. Statistical indistinguishability and absolute structural consistency within these nodes may reflect an early anatomical locus of shared vulnerability, consistent with the NES seeding stage [8] (Figure 6, Table 3).

Morphological symmetry in the *LH Transverse Temporal* cortex may be compatible with alterations in neuromodulatory systems (e.g., noradrenergic and serotonergic pathways), although this interpretation cannot be directly tested with the present data. As the site of the primary auditory cortex (Heschl’s gyrus), this region relies on dense projections from the *Locus Coeruleus* and the *Raphe Nuclei* to maintain cortical arousal and auditory signal-to-noise modulation [8, 48–50]. Volumetric analysis identifies a hemispheric gradient within this manifold: significant structural decay characterizes the right hemisphere (*p* = 0.0025), while the left remains statistically contiguous with the healthy baseline (*p* = 0.0606). This lateralized vulnerability suggests that sensory hub degradation precedes the collapse of associative networks.

Simultaneously, bilateral *Choroid Plexus* hypertrophy is consistent with the possibility of altered system-level homeostatic regulation. As the primary site for cerebrospinal fluid (CSF) production and glymphatic waste clearance, invariant Choroid Plexus expansion suggests that impaired proteostatic drainage precedes the accumulation of disease-specific aggregates [51, 52]. The *RH Choroid Plexus* (*p* = 0.051) represents a node of maximum convergence within a context of brain-wide divergence. While the magnitude of this hit is more pronounced in AD (*p <* 0.001), consistent with [53], the underlying attempt to manage proteinopathic overload through increased CSF production remains synchronized across phenotypes.

### Molecular Correlates of Structural Anchors

Master anchor identification via IIT logic converges with recent large-scale plasma proteomics delineating pan-neurodegenerative molecular signatures [11]. Synchronized volumetric expansion of the *RH Choroid Plexus* serves as a macrostructural correlate to shared dysregulation in proteins governing extracellular matrix (ECM) remodeling and immune response. Specifically, the consistent elevation of SMOC1 and the enrichment of ECM organization pathways—including multiple matrix metalloproteinases—across AD and PD cohorts [11] point toward systemic alterations at cerebral homeostatic interfaces. As a critical blood-CSF barrier, the Choroid Plexus undergoes structural remodeling in response to these shared proteinopathic hits, mirroring a universal collapse in proteostasis and glymphatic clearance that precedes clinical bifurcation.

Similarly, the structural invariance of the *LH Transverse Temporal* cortex aligns with identified synaptic and innate immune disruptions common to both disorders. Pan-neurodegenerative downregulation of the synaptic protein NPTXR and dysregulation of MAPK1-mediated immune signaling [11] substantiate a synchronized pathological seeding phase. Within this framework, the Transverse Temporal master anchor identifies a neuroanatomical “ground zero” where shared molecular failures manifest as stable structural patterns. These signatures persist until phenotype-specific cascades—such as apoptotic pathways in AD or ubiquitin-proteasome impairments in PD—drive phenotypic bifurcation into distinct clinical silos. By prioritizing these structural anchors, IIT effectively distills the neuroanatomical manifestation of the pan-neurodegenerative proteome.

### Parkinson’s Disease as a Morphological Transition State

Volumetric distributions identify PD as a *transitional state* within the neurodegenerative continuum. Across both Master Anchors and Transition Pillars, the PD cohort consistently occupies an intermediate position between CN and the AD group. The PD brain exhibits incipient signs of the shared pathological trajectory without reaching the systemic collapse of advanced AD. IIT captures this latent footprint before it becomes statistically overt through standard frequentist testing (Figure 6b, *ns* vs. Healthy).

The residence of PD volumes within a statistical “gray zone”—not significantly differentiated from AD yet contiguous with health—is consistent with the chronological window of the NES seeding phase. This homeostatic hit, centered on the Choroid Plexus expansion, is supported by the “expansion model” [43], suggesting that regional enlargement reflects a compensatory but failing effort to wash out metabolic toxins rather than the volumetric reduction reported in other studies [54]. In this framework, the non-significant difference between PD and CN highlights the clinical bifurcation, where a shared homeostatic struggle evolves into phenotype-specific decay.

### Limbic Bifurcation and Hybrid Pathological Nodes

The common neuroanatomical trunk eventually yields to phenotype-specific atrophy. During bifurcation, differential regional kinetics of *α*-synuclein and tau proteinopathies drive the structural collapse of limbic and temporal networks, yielding the pronounced statistical separation observed in the *Hippocampus* and *Amygdala* [24, 55]. High effect sizes mark the threshold where proteinopathic loads eclipse shared monoaminergic failure.

Within this landscape, the *RH Amygdala* functions as a critical hybrid node. While its involvement in both pathologies identifies it as a structural anchor, its high discriminative rank reveals a degradation morphology that remains pathology-dependent. Its high connectivity and metabolic cost render it a primary site for differential atrophy, acting as a bridge between shared vulnerability and diagnostic divergence.

The negligible Effect Size (0.2860) in the *LH Transverse Temporal* master anchor introduces a distinction between atrophy markers and backbone anchors. While traditional paradigms discard this region due to low pathological impact, IIT identifies it as a shared “control node” and a structural bridge of stability. This sensory ground zero may reflect the synchronized denervation of norepinephrine and serotonin, establishing a common neuroanatomical footprint before clinical bifurcation into cognitive or motor silos [8, 56–58].

### 3.4 Etiological Drivers of the Neuroanatomical Backbone

Identification of the *RH Choroid Plexus* and *LH Transverse Temporal* cortex as master anchors aligns with the shared risk profile characteristic of the NES seeding stage [8]. Common environmental triggers, including high iron intake and pesticide exposure, drive this invariant structural backbone by inducing early homeostatic failure and proteostatic stress [59, 60]. Specifically, expansion of the choroid plexus across both cohorts suggests a shared response to impaired waste clearance, potentially exacerbated by systemic factors such as high cholesterol, which compromise blood-CSF barrier integrity in both phenotypes [61].

Conversely, etiological factors with divergent risk profiles modulate the transition to the bifurcation stage—captured through the inductive assessment of discriminative features. Agents such as nicotine and coffee exert antagonistic effects, offering protection for the PD trajectory while potentially increasing vulnerability in the AD path [8, 62]. This etiological divergence mirrors the behavior of the identified *Transition Pillars* (e.g., Amygdala and Hippocampus), which maintain absolute rank-order consistency (*ρ* = 1.0) despite significant divergence in atrophy magnitude. These findings suggest the possibility that while shared genetic and environmental factors may contribute to an invariant neuroanatomical core, subsequent interactions with lifestyle-related variables could play a role in the bifurcation into divergent clinical manifestations. This is consistent with the NES hypothesis that AD and PD potentially share early structural features before diverging.

## 4 Conclusion

This study introduces a framework for identifying candidate shared neuroanatomical features across neurodegenerative conditions. Implementation of IIT logic suggests that a stable set of structural anchors remains invariant across the neurodegenerative spectrum, notwithstanding divergent symptomatic expressions. Identification of this unified backbone yields quantitative preliminary empirical support for the NES hypothesis, facilitating a perspective shift from phenotype-specific silos toward a holistic, system-level view on brain aging [19].

These insights carry profound implications for geriatric neurology and biomarker discovery. Isolation of the invariant pathological core pinpoint anatomical landmarks critical to the subclinical seeding and compensatory stages of neurodegeneration. By shifting the analytical objective from maximal group separation to structural commonality, this methodology provides a robust foundation for early intervention and cross-disciplinary knowledge propagation. Ultimately, this approach transforms ML from a descriptive classifier into a discovery engine capable of elucidating universal structural laws governing complex neurodegenerative processes.

While the identified backbone appears robust, several limitations should be considered. First, although the analysis incorporates longitudinal information through the Cumulative Structural Integrity Index (CSII), this aggregation provides a temporally averaged representation rather than explicitly modeling disease dynamics, thus limiting inference on progression trajectories. Second, the study is based on two cohorts (ADNI and PPMI), and the lack of external validation may constrain the generalizability of the findings. Finally, structural MRI alone cannot resolve the underlying molecular mechanisms. Although the identified backbone is consistent with hypothesized monoaminergic and glymphatic dysfunction, direct validation through molecular biomarkers—such as *α*-synuclein and tau PET imaging—remains necessary to substantiate the proposed proteinopathic drivers. The integration of multimodal data, including proteomic and metabolomic profiles, within this framework may further refine the characterization of structural anchors and support the development of more precise, personalized approaches to neurodegenerative aging [63, 64].

## 5 Methods

### 5.1 Cohort Selection and Data Acquisition

Data originate from two large-scale repositories: the Alzheimer’s Disease Neuroimaging Initiative (ADNI)^1^ and the Parkinson’s Progression Markers Initiative (PPMI)^2^. ADNI tracks neurodegenerative progression from mild cognitive impairment to AD through integrated longitudinal markers, while PPMI comprises a comprehensive multi-modal dataset spanning idiopathic and genetic PD cases alongside CN.

The finalized cohort includes 303 individuals distributed equally across three experimental groups: Healthy Controls (CN, *n* = 101), AD (*n* = 101), and PD (*n* = 101). Strict matching for age, sex, and total intracranial volume ensures cohort comparability and minimizes demographic confounding.

### 5.2 MRI Acquisition and Image Processing

High-resolution T1-weighted sequences provide the foundation for precise morphometric assessment. The dataset utilizes two optimized protocols: *MPRAGE* and *MPRAGE GRAPPA*, which leverages parallel imaging to reduce scan time without compromising image fidelity [65, 66]. Technical specifications across platforms are standardized to ensure cross-cohort homogeneity (Table 4).

**Table 4.** Standardized acquisition parameters for T1-weighted scans.

| Parameter | ADNI (Trio) | PPMI (Verio) |
| --- | --- | --- |
| Sequence | MPRAGE | MPRAGE |
| Acquisition plane | Sagittal (3D) | Sagittal (3D) |
| TR / TE / TI (ms) | 2300 / 3.0 / 900 | 2300 / 3.0 / 900 |
| Flip angle (°) | 9 | 9 |
| Slice thickness (mm) | 1.2 | 1.0 |
| Matrix size | 240 × 256 × 160 | 240 × 256 × 176 |
| Pixel spacing (mm) | 1.0 × 1.0 | 1.0 × 1.0 |

**Table 5.** Regional volumetric characterization of the neurodegenerative manifold. Detailed mapping of mean volumes and atrophy percentages across Healthy Controls (CN), Parkinson’s disease (PD), and Alzheimer’s disease (AD) cohorts. Regional volumes are expressed in ×10^3^ mm^3^. *PD*_A_ and *AD*_A_ represent percentage atrophy relative to the healthy baseline; negative values identify regions of volumetric expansion. Pairwise *p*-values (*p_CN/PD_* and *p_CN/AD_*) indicate statistical significance relative to the control group.

| Region | CN | PD | AD | $PD_A$ | $AD_A$ | $p_{CN/PD}$ | $p_{CN/AD}$ |
| --- | --- | --- | --- | --- | --- | --- | --- |
| *Subcortical, Limbic* |  |  |  |  |  |  |  |
| Brain Stem | 20.85 | 21.95 | 19.78 | −5.25 | 5.15 | 0.03 | 0.00 |
| LH Accum | 0.49 | 0.50 | 0.38 | −2.79 | 21.77 | 1.00 | 0.00 |
| LH Amyg | 1.67 | 1.72 | 1.14 | −2.82 | 32.21 | 0.59 | 0.00 |
| LH Caud | 3.44 | 3.47 | 3.31 | −0.69 | 3.87 | 1.00 | 0.00 |
| LH Cereb Cortex | 50.81 | 52.71 | 47.07 | −3.74 | 7.36 | 0.11 | 0.00 |
| LH Cereb WM | 14.00 | 14.82 | 13.12 | −5.86 | 6.28 | 0.04 | 0.00 |
| LH Ent | 1.75 | 1.82 | 1.15 | −4.11 | 34.11 | 0.56 | 0.00 |
| LH Hipp | 4.02 | 4.09 | 2.88 | −1.88 | 28.36 | 1.00 | 0.00 |
| LH Pall | 2.00 | 2.08 | 1.88 | −4.09 | 6.13 | 0.14 | 0.00 |
| LH Put | 4.62 | 4.61 | 4.19 | 0.25 | 9.25 | 1.00 | 0.00 |
| LH Thal | 7.00 | 7.32 | 6.07 | −4.56 | 13.30 | 0.04 | 0.00 |
| RH Accum | 0.50 | 0.52 | 0.39 | −3.54 | 21.83 | 1.00 | 0.00 |
| RH Amyg | 1.78 | 1.88 | 1.26 | −5.51 | 28.90 | 0.06 | 0.00 |
| RH Caud | 3.63 | 3.68 | 3.44 | −1.33 | 5.18 | 1.00 | 0.00 |
| RH Cereb Cortex | 51.80 | 53.71 | 47.59 | −3.69 | 8.12 | 0.16 | 0.00 |
| RH Cereb WM | 13.35 | 14.16 | 12.57 | −6.03 | 5.84 | 0.02 | 0.00 |
| RH Ent | 1.72 | 1.76 | 1.12 | −2.35 | 35.16 | 1.00 | 0.00 |
| RH Hipp | 4.09 | 4.22 | 2.99 | −3.13 | 26.82 | 0.40 | 0.00 |
| RH Pall | 1.91 | 2.01 | 1.84 | −5.20 | 3.90 | 0.03 | 0.00 |
| RH Put | 4.67 | 4.68 | 4.29 | −0.23 | 8.15 | 1.00 | 0.00 |
| RH Thal | 6.89 | 7.18 | 6.02 | −4.15 | 12.64 | 0.05 | 0.00 |
| *Cortical (LH)* |  |  |  |  |  |  |  |
| LH Cau Ant Cing | 2.72 | 2.94 | 2.54 | −8.09 | 6.64 | 0.01 | 0.00 |
| LH Cau Mid Front | 5.63 | 5.89 | 5.01 | −4.72 | 10.98 | 0.31 | 0.00 |

Table 5 – Continued from previous page
| Region | CN | PD | AD | $PD_{\mathbf{A}}$ | $AD_{\mathbf{A}}$ | $p_{CN/PD}$ | $p_{CN/AD}$ |
| --- | --- | --- | --- | --- | --- | --- | --- |
| LH Cuneus | 4.12 | 4.27 | 3.72 | -3.59 | 9.61 | 0.54 | 0.00 |
| LH Fusiform | 7.47 | 7.79 | 6.18 | -4.22 | 17.26 | 0.29 | 0.00 |
| LH Inf Pariet | 10.61 | 11.01 | 9.26 | -3.81 | 12.72 | 0.46 | 0.00 |
| LH Inftemp | 10.82 | 11.17 | 8.56 | -3.23 | 20.90 | 0.78 | 0.00 |
| LH Insula | 6.39 | 6.61 | 5.57 | -3.49 | 12.79 | 0.51 | 0.00 |
| LH Lat Occip | 9.62 | 9.93 | 8.90 | -3.21 | 7.53 | 0.55 | 0.00 |
| LH Lat Orbitofront | 4.79 | 5.04 | 4.36 | -5.18 | 9.06 | 0.09 | 0.00 |
| LH Med Orbitofront | 4.09 | 4.27 | 3.73 | -4.42 | 8.77 | 0.20 | 0.00 |
| LH Midtemp | 11.60 | 12.15 | 9.34 | -4.73 | 19.49 | 0.15 | 0.00 |
| LH Paracen | 3.46 | 3.61 | 3.17 | -4.36 | 8.38 | 0.23 | 0.00 |
| LH Parahipp | 2.65 | 2.77 | 2.04 | -4.46 | 23.11 | 0.25 | 0.00 |
| LH Pars Orbit | 1.81 | 1.92 | 1.71 | -5.88 | 5.30 | 0.07 | 0.00 |
| LH Pars Operc | 3.41 | 3.55 | 3.08 | -4.18 | 9.64 | 0.28 | 0.00 |
| LH Pars Triang | 3.44 | 3.62 | 3.31 | -5.14 | 3.79 | 0.25 | 0.00 |
| LH Pericalc | 1.78 | 1.84 | 1.62 | -3.35 | 8.79 | 0.64 | 0.00 |
| LH Poscen | 9.33 | 9.70 | 8.63 | -4.03 | 7.51 | 0.08 | 0.00 |
| LH Post Cing | 3.00 | 3.14 | 2.64 | -4.59 | 12.07 | 0.13 | 0.00 |
| LH Precen | 10.15 | 10.53 | 9.38 | -3.76 | 7.59 | 0.39 | 0.00 |
| LH Precun | 8.20 | 8.87 | 6.91 | -8.23 | 15.70 | 0.01 | 0.00 |
| LH Rost Ant Cing | 2.47 | 2.64 | 2.21 | -6.69 | 10.61 | 0.03 | 0.00 |
| LH Rost Mid Front | 14.52 | 15.16 | 12.98 | -4.42 | 10.64 | 0.21 | 0.00 |
| LH Sup Front | 14.02 | 14.62 | 12.64 | -4.31 | 9.82 | 0.21 | 0.00 |
| LH Sup Pariet | 8.83 | 9.21 | 7.80 | -4.29 | 11.69 | 0.29 | 0.00 |
| LH Sup Temp | 8.79 | 9.19 | 7.65 | -4.56 | 12.93 | 0.21 | 0.00 |
| LH Supramarg | 7.40 | 7.70 | 6.54 | -4.01 | 11.61 | 0.31 | 0.00 |
| LH Transv Temp | 1.68 | 1.75 | 1.48 | -4.11 | 11.74 | 0.31 | 0.00 |
| *Cortical (RH)* |  |  |  |  |  |  |  |
| RH Cau Ant Cing | 2.57 | 2.75 | 2.36 | -7.01 | 8.23 | 0.01 | 0.00 |
| RH Cau Mid Front | 5.14 | 5.38 | 4.59 | -4.72 | 10.68 | 0.27 | 0.00 |
| RH Cuneus | 4.03 | 4.17 | 3.65 | -3.53 | 9.49 | 0.56 | 0.00 |
| RH Fusiform | 7.66 | 7.96 | 6.32 | -3.94 | 17.47 | 0.34 | 0.00 |
| RH Inf Pariet | 10.39 | 10.81 | 9.20 | -4.07 | 11.44 | 0.29 | 0.00 |
| RH Inftemp | 10.90 | 11.27 | 8.62 | -3.39 | 20.88 | 0.56 | 0.00 |
| RH Insula | 6.28 | 6.51 | 5.44 | -3.64 | 13.43 | 0.46 | 0.00 |
| RH Lat Occip | 9.48 | 9.79 | 8.70 | -3.29 | 8.21 | 0.52 | 0.00 |
| RH Lat Orbitofront | 4.83 | 5.06 | 4.38 | -4.76 | 9.34 | 0.12 | 0.00 |
| RH Med Orbitofront | 4.11 | 4.28 | 3.75 | -4.07 | 8.68 | 0.25 | 0.00 |
| RH Midtemp | 11.73 | 12.52 | 9.71 | -6.72 | 17.21 | 0.01 | 0.00 |
| RH Paracen | 3.35 | 3.48 | 3.10 | -3.88 | 7.50 | 0.36 | 0.00 |
| RH Parahipp | 2.67 | 2.80 | 2.09 | -4.71 | 21.79 | 0.24 | 0.00 |
| RH Pars Operc | 3.38 | 3.52 | 3.06 | -4.04 | 9.43 | 0.30 | 0.00 |
| RH Pars Orbit | 1.86 | 2.01 | 1.83 | -7.83 | 1.63 | 0.00 | 0.00 |
| RH Pars Triang | 3.39 | 3.47 | 3.16 | -2.29 | 6.86 | 1.00 | 0.00 |
| RH Pericalc | 1.72 | 1.77 | 1.56 | -2.94 | 9.30 | 0.71 | 0.00 |
| RH Poscen | 8.65 | 8.93 | 8.03 | -3.18 | 7.22 | 0.16 | 0.00 |
| RH Post Cing | 3.08 | 3.23 | 2.69 | -4.90 | 12.71 | 0.12 | 0.00 |
| RH Precen | 10.26 | 10.63 | 9.45 | -3.63 | 7.86 | 0.41 | 0.00 |
| RH Precun | 8.85 | 9.50 | 7.57 | -7.35 | 14.40 | 0.01 | 0.00 |
| RH Rost Ant Cing | 2.52 | 2.66 | 2.23 | -5.42 | 11.47 | 0.06 | 0.00 |
| RH Rost Mid Front | 14.01 | 14.56 | 12.56 | -3.92 | 10.36 | 0.28 | 0.00 |
| RH Sup Front | 13.62 | 14.16 | 12.25 | -3.96 | 10.05 | 0.27 | 0.00 |

Table 5 – Continued from previous page
| Region | CN | PD | AD | $PD_{\mathbf{A}}$ | $AD_{\mathbf{A}}$ | $PCN/PD$ | $PCN/AD$ |
| --- | --- | --- | --- | --- | --- | --- | --- |
| RH Sup Pariet | 8.69 | 9.03 | 7.72 | -3.95 | 11.16 | 0.36 | 0.00 |
| RH Sup Temp | 8.91 | 9.29 | 7.70 | -4.28 | 13.55 | 0.24 | 0.00 |
| RH Supramarg | 7.23 | 7.51 | 6.39 | -3.88 | 11.63 | 0.33 | 0.00 |
| RH Transv Temp | 1.65 | 1.71 | 1.46 | -3.88 | 11.52 | 0.33 | 0.00 |
| *White Matt, Ventr, Flui* |  |  |  |  |  |  |  |
| CSF | 1.15 | 1.31 | 1.54 | -14.38 | -34.07 | 0.00 | 0.00 |
| LH Cerebr White Mat | 221.23 | 232.12 | 199.60 | -4.92 | 9.78 | 0.05 | 0.00 |
| LH Chor Plex | 0.79 | 0.87 | 1.00 | -10.40 | -26.08 | 0.06 | 0.00 |
| LH Lat Vent | 13.28 | 15.72 | 24.93 | -18.37 | -87.78 | 0.23 | 0.00 |
| RH Cerebr White Mat | 221.08 | 232.29 | 200.39 | -5.07 | 9.36 | 0.03 | 0.00 |
| RH Chor Plex | 0.86 | 0.93 | 1.02 | -7.69 | -18.17 | 0.46 | 0.00 |
| RH Lat Vent | 11.97 | 14.09 | 22.18 | -17.72 | -85.28 | 0.42 | 0.00 |

The FastSurfer pipeline, a GPU-accelerated implementation of FreeSurfer, executes automated segmentation and parcellation [66]. This framework integrates HypVINN and CerebNet modules to extend traditional parcellation to hypothalamic and cerebellar structures [67, 68]. Preprocessing incorporates motion correction, intensity normalization, and affine registration to the MNI305 coordinate space, followed by automated skull stripping and subcortical segmentation via atlas-based probabilistic models. Spherical registration maps cortical folding patterns to an atlas template, yielding a robust feature set encompassing gray matter volume, cortical thickness, and surface area across 84 regions.

To provide a proxy of cumulative structural burden, the Cumulative Structural Integrity Index (CSII) aggregates longitudinal volumes across three time points (0, 12, and 24 months): *CSII* = (*V*_0_ + *V*_12_ + *V*_24_)*/*3. This temporal integration minimizes measurement noise and optimizes the signal-to-noise ratio for subsequent analysis. It smooths temporal variability but does not explicitly model rates of change or disease trajectories. The analytical framework evaluates the complete set of 84 neuroanatomical descriptors to provide a comprehensive mapping of the neurodegenerative manifold. Statistical models incorporate sex as a biological variable to account for potential dimorphic patterns of decay across the shared pathological core, ensuring that identified structural anchors remain invariant regardless of biological sex.

### 5.3 Inductive Benchmarking and Model Generalization

The evaluation of diagnostic robustness follows a dual-path strategy to bifurcate model benchmarking from structural discovery. The inductive path establishes clinical reliability through three optimized architectures: *Random Forest* (RF), *Gradient Boosting* (GB), and *Logistic Regression* (LR). This selection encompasses both a regularized linear baseline (LR) and models capable of capturing high-order non-linear interactions (RF, GB).

A stratified 5-fold cross-validation scheme, repeated across 10 independent stochastic realizations, ensures that performance metrics remain uninflated by data leakage. Within each loop, all preprocessing—including median imputation for missing volumes and *Z*-score standardization—is performed nested within the training fold. An automated *GridSearchCV* protocol executes hyperparameter optimization within these folds: RF utilizes 500 estimators with balanced class weights; GB incorporates stochastic subsampling (0.8); and LR employs an *L*_2_ penalty with optimized *C* coefficients. Evaluation prioritizes ROC-AUC, Accuracy, and *F* 1-macro to capture decision-boundary sensitivity and stability under cohort imbalance.

### 5.4 Transductive Discovery via the IIT Framework

While benchmarking ensures reliability, identification of shared structural principles follows a transductive, unsupervised domain adaptation logic. Unlike the inductive path, the IIT calculation leverages the entire disease-specific manifold (AD and PD) to maximize statistical resolution and identify common pathological laws.

This process first estimates regional importance via a consensus-based Borda score (*B_i_*) [69, 70]. Optimized RF, GB, and LR architectures undergo training on the full cohort, and the average importance rank across these *M* = 3 models defines the score: 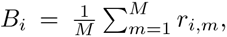 where *r_i,m_* denotes the rank (e.g., *r* = 1 for the most discriminative). Low *B_i_* values identify “Bifurcation Markers” responsible for structural separation, while high *B_i_* values signify discriminative neutrality and serve as candidates for stable structural anchors.

The IIT mechanism [17] then calculates a directed Importance Inversion Transfer Score 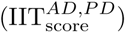 by integrating Borda neutrality with metric consistency (Spearman’s rank correlation *ρ*) and geometric proximity (normalized mean difference Δ):

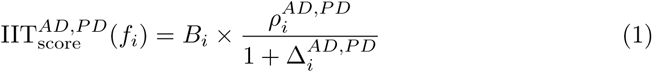

where 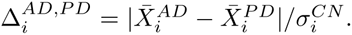 Spearman’s *ρ* ensures robustness against biological outliers and prioritizes rank-order consistency. The denominator functions as an inverse mean difference penalty, ensuring that selected anchors identify pathological hits preceding symptomatic bifurcation. The framework selects the ten descriptors with the highest scores as the primary NES core anchors.

### 5.5 Dimensionality Control and Statistical Validation

The reduction of 84 initial volumetric descriptors into ten stable invariants optimizes the effective subject-to-feature ratio from *N/p ≈* 3 to *N/p ≈* 30. This dimensionality control ensures that performance gains reflect robust pathological principles rather than numerical artifacts. Non-parametric statistical validation utilizes the Kruskal-Wallis *H*-test to assess global variance, while Dunn’s post-hoc analysis with Bonferroni correction provides the framework for multiple comparisons. Formal identification of structural anchors relies on the criterion of statistical indistinguishability between AD and PD cohorts (*p_AD/P D_ ≥* 0.05), with observed trends further substantiated through paired two-tailed *t*-tests.

### 5.6 Computational Infrastructure

The framework integrates scikit-learn for ML and PyTorch for tensor operations. A dynamic GPU-selection logic (utilizing nvidia-smi) assigns computations to the most efficient resource (NVIDIA RTX 4090 or A100-PCIE). Fixed 10 independent random seeds across all modules guarantee computational determinism. CUDA-optimized operations within PyTorch reduce empirical estimation latency by over one order of magnitude compared to CPU-based implementations.

## 6 Supplementary Information

### 6.1 Exhaustive Volumetric Profiling of the Neurodegenerative Manifold

This supplementary section presents a systematic topographical mapping of regional brain volumes across 84 neuroanatomical regions. Bootstrapped ensemble means characterize the CN, PD, and AD cohorts. Kruskal-Wallis H-tests evaluated global variance, while pairwise post-hoc Dunn’s tests with Bonferroni correction adjusted for multiple comparisons. Analysis supports a pervasive structural impact in AD, involving approximately 87% of the investigated manifold, in contrast to the more localized, region-specific alterations characterizing PD. Negative atrophy values denote volumetric expansion relative to the healthy baseline, identifying regions potentially subject to neuro-inflammatory or compensatory mechanisms.

## Acknowledgements.

This work was supported by the National Research Council (CNR), Italy. The authors acknowledge the Institute of Cognitive Sciences and Technologies (ISTC-CNR) for providing the necessary research infrastructure.

Data used in the preparation of this article were obtained from the Alzheimer’s Disease Neuroimaging Initiative (ADNI) database (adni.loni.usc.edu) and the Parkinson’s Progression Markers Initiative (PPMI) database (www.ppmi-info.org/data). As such, the investigators within the ADNI and PPMI contributed to the design and implementation of their respective studies and/or provided data but did not participate in the analysis or writing of this report.

ADNI data collection and sharing was funded by the Alzheimer’s Disease Neuroimaging Initiative (National Institutes of Health Grant U01 AG024904) and DOD ADNI (Department of Defense award number W81XWH-12-2-0012). PPMI – a public-private partnership – is funded by the Michael J. Fox Foundation for Parkinson’s Research and funding partners (a full list of partners is available at https://www.ppmi-info.org/about-ppmi/who-we-are/study-sponsors).

## Author Contributions

D.C. conceived the study and developed the IIT mechanism. Author contributions according to the CRediT taxonomy are as follows: D.C.: Conceptualization, Methodology, Software, Validation, Formal analysis, Investigation, Resources, Data curation, Writing – original draft, Writing – review & editing, Visualization, Supervision, Project administration, and Funding acquisition. S.T.: Methodology, Software, Validation, Resources, and Data curation. All authors read and approved the final manuscript.

## Ethics approval and consent to participate

This study utilized anonymized secondary data from the Alzheimer’s Disease Neuroimaging Initiative (ADNI) and the Parkinson’s Progression Markers Initiative (PPMI). Ethical approval and informed consent were obtained by the original data collection initiatives. All procedures were conducted in accordance with the relevant data use agreements and ethical guidelines.

## Consent for publication

Not applicable. This manuscript does not contain any identifying information or individual person’s data in any form.

## Data Availability

The neuroimaging data analyzed in this study were obtained from the Alzheimer’s Disease Neuroimaging Initiative (ADNI) (adni.loni.usc.edu) and the Parkinson’s Progression Markers Initiative (PPMI) (ppmi-info.org). These datasets are available to researchers upon registration and approval of a data utility request through the respective consortium portals.

## Materials availability

Not applicable. No physical materials were generated or used in this study.

## Code Availability

Custom Python code implementing the IIT mechanism and the complete statistical validation suite is available as a comprehensive Jupyter Notebook on GitHub: https://github.com/danielecaligiore/IIT NES.

## Footnotes

1 adni.loni.usc.edu

2 ppmi-info.org

